# Timed collinear activation of Hox genes during gastrulation controls the avian forelimb position

**DOI:** 10.1101/351106

**Authors:** Chloe Moreau, Paolo Caldarelli, Didier Rocancourt, Julian Roussel, Nicolas Denans, Olivier Pourquie, Jerome Gros

## Abstract

Limb position along the body is highly consistent within one species but very variable among vertebrates. Despite major advances in our understanding of limb patterning in three dimensions, how limbs reproducibly form along the anteroposterior axis remains largely unknown. Hox genes have long been suspected to control limb position, however supporting evidences are mostly correlative and their role in this process remains unclear. Here we show that Hox genes determine the avian forelimb position in a two-step process: first, their sequential collinear activation during gastrulation controls the relative position of their own successive expression domains along the body axis. Then, within these collinear domains, Hox genes differentially activate or repress the genetic cascade responsible for forelimb initiation. Furthermore, we provide evidences that changes in the timing of collinear Hox gene activation might underlie natural variation in forelimb position between different birds. Altogether our results which characterize the cellular and molecular mechanisms underlying the regulation and natural variation of forelimb position in avians, show a direct and early role for Hox genes in this process.

## Introduction

In tetrapods, limbs are always positioned at the level of the cervico-thoracic (forelimb) or lumbo-sacral (hindlimb) vertebral transitions, however the position of these frontiers varies greatly among species. Almost all mammals form forelimbs at the level of the 8^th^ vertebrae, birds display tremendous variation with sparrow, chicken and swans forming forelimbs at the level of the 10^th^, 15^th^ and 25^th^ vertebrae, respectively. Frogs, in turn, exhibit forelimbs at the level of the 2^nd^ vertebrae. Hox genes have long been proposed to regulate limb position during development (for review see [1–3]). Organized in four different clusters, they display a chromosomal organization that reflects their sequential timing of expression and their successive domains of expression along the antero-posterior (A-P) axis, named temporal and spatial collinearity, respectively [4–6]. The observation that anterior boundaries of expression of specific Hox genes match forelimb, interlimb and hindlimb borders across tetrapod species [7–9] and more recently that Hox genes can bind an enhancer of Tbx5, a transcription factor essential for forelimb initiation [10,11], led to the proposition that Hox genes might regulate limb position. However, whereas gain and loss of function of single or multiple Hox genes result in the transformation of vertebral identity [12], none of the reported Hox mutants in mice shows a major phenotype regarding limb position. Therefore, whether Hox genes control limb initiation and positioning has not been established.

Limbs originate from the somatopleural Lateral Plate Mesoderm (LPM), an epithelial layer of tissue which flanks axial embryonic structures. Surprisingly, whereas the cellular events underlying the formation of embryonic compartments adjacent to the LPM are well described (i.e. neural tube, notochord, somites, and intermediate mesoderm)[13–16], little is known about how the LPM is generated during gastrulation. Lineage tracing experiments in the chick have revealed that LPM precursor cells arise from the central third of the primitive streak [15–19]. Only one lineage analysis, using colored chalk dust [20], traced back the origin of forelimb, interlimb and hindlimb fields up to stage 7 (i.e. after LPM cells have initiated their ingression through the primitive streak). A series of grafting studies demonstrated that the forelimb identity becomes determined by at least stage 9 [21] and that the forelimb field carries the morphogenetic potential to induce a limb in the flank by stage 11 [22]. However, such studies were all designed an interpreted with respect to the determination of limb field identity (forelimb vs. hindlimb) [21], polarity (antero-posterior or dorso-ventral) [23] or morphogenetic inductive potential [22,24,25] and did not address the limb- vs. non-limb-forming domains determination and therefore did not address limb position determination. Thus, how and even when the LPM becomes determined into limb and non-limb forming domains remains to be investigated.

Here, we examine how the forelimbs are positioned along the A-P axis during avian development. We find that the LPM is progressively formed and concomitantly patterned by Hox genes into limb- and non-limb forming domains, during the process of gastrulation. Moreover we provide data suggesting that relative changes in the timing of Hox collinear activation might underlie natural variation in forelimb position observed between different bird species.

## Results

### Forelimb position is already determined by the end of gastrulation

We first sought to elucidate when the position of the future forelimb is determined. The LPM which is generated during gastrulation epithelializes into the somatopleure by stage 11 (i.e. 2 days of development). To test whether limb position is already determined in the freshly generated LPM, we micro-dissected, rotated and grafted back the right somatopleure encompassing both forelimb and interlimb prospective domains of stage 11 chicken embryos (Figure 1A). Operated embryos were re-incubated for 48h, until limbs have clearly formed. 65% of operated embryos exhibited either a total or partial posterior shift of the forelimb, as revealed by expression of the limb marker FGF10 (Figures 1B and 1C, n=15/23). Shifted limb buds also expressed Tbx5, demonstrating that the position of the forelimb field has been displaced by the surgical procedure (Figure 1D, n=2/2). To rule out that the shift of the limb bud territory resulted from artificial induction of donor surrounding tissues, we performed quail-chick grafting experiments to verify that the grafted tissue contained only somatopleural cells. To directly visualize the grafted tissue, we generated a transgenic quail line expressing membrane-bound eGFP under the control of the ubiquitous human Ubiquitin C (hUbC) promoter (*hUbC:memGFP*), (Figure 1E). Transverse sections of grafted embryos clearly show that only the somatopleural LPM (GFP positive) was transplanted at the level of forelimb or interlimb (Figures 1F-1I). Altogether these experiments show that the position of the forelimb domain relative to the interlimb domain is established autonomously within the somatopleural LPM and as early as stage 11.

**Figure 1.**
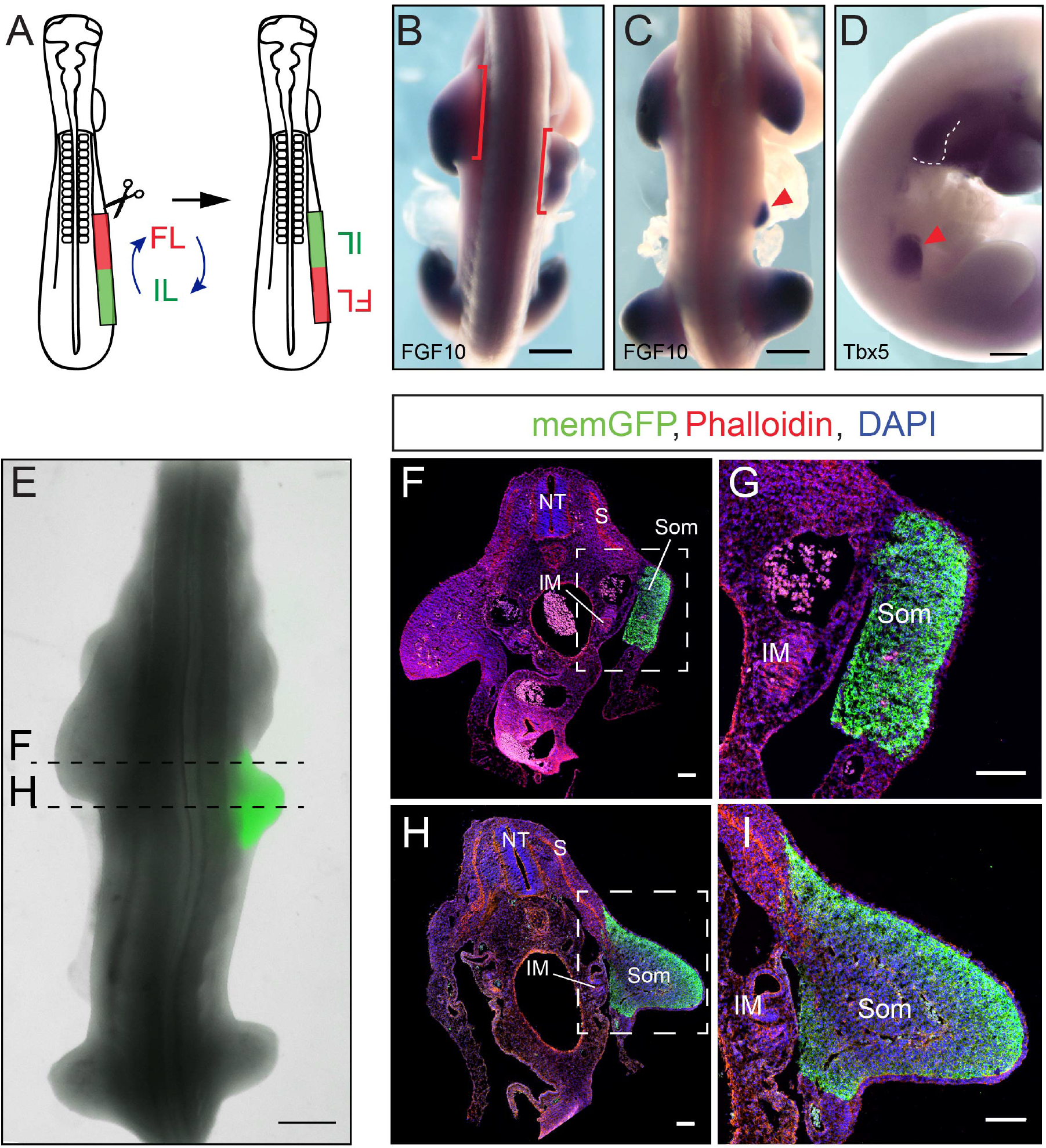
Forelimb position is already determined after gastrulation. (A) Representation of the transplantation experiment in a stage 11 chicken embryo. FL, Forelimb; Il, interlimb. (B-D) FGF10 (B-C) and Tbx5 (D) expression 48h after transplantation experiment, showing complete (B, n=3/23 embryos, Red brackets) or partial (C, D, n=12/23 embryos, red arrowhead) displacement of the forelimb bud. (E) Chicken embryo grafted with memGFP transgenic quail somatopleure (green) showing a posterior displacement of the forelimb (n=3/3 embryos). (F-I) Transverse sections of quail-chick chimera at the forelimb (F-G) and interlimb (H-I) levels, stained with phalloidin (red), GFP antibody (green), and DAPI (blue). (G) and (I) are higher magnifications of (F) and (H), respectively. NT, neural tube; S, somite; IM, intermediate mesoderm; Som, somatopleure. Scale bar is 500Sm in (B-E), and 100μm in (F-I).

### Forelimb, interlimb and hindlimb fields are sequentially formed during gastrulation

The finding that the LPM is patterned into forelimb and interlimb domains by stage 11 raises the possibility that it might be patterned even earlier. The origin of the LPM in the epiblast has been traced back to the middle third of the primitive streak (PS) in chicken embryos [15,16,19], however how forelimb, interlimb and hindlimb cells are specifically generated has not been characterized. We performed a dynamic lineage analysis of prospective limb precursor cells by electroporating fluorescent reporters (GFP and nuclear H2b-RFP) into the presumptive LPM territory of stage 4 chicken embryos. Electroporated embryos were then cultured ex-vivo [26] and imaged using 2-photon video-microscopy for approximately 24h (Figure 2A and Movie S1). Retrospective tracking of LPM precursors identified the epiblast origin of forelimb, interlimb and hindlimb fields (Movie S2). First, we observed that the formation of the LPM takes place between stage 4 and 10 (i.e. spanning 24h of development). Second, we could determine that most forelimb precursor cells are generated between stage 4 and 5, whereas interlimb and hindlimb precursor cells are gradually generated at later stages, mostly at stage 6-7 and 8-9, respectively (Figure 2B). To confirm these results using a different lineage tracing technique, we generated a transgenic quail line expressing the Green-to-Red photoconvertible fluorescent protein mEOS2 under the control of the ubiquitous hUbC promoter (*hUbC:mEOS2FP*). In *hUbC:mEOS2FP* transgenic quail embryos, regions could be very precisely photoconverted, and readily tracked for 24h. We thus photoconverted the prospective LPM cells in the PS of stage 6, 7, 9 and 10 mEOS2 transgenic embryos (Figures S1A-S1D) and followed their fate for 24h. As expected, the later the cells were photoconverted in the primitive streak, the more posterior they localized in the LPM (Figures S1A’-S1D’, S1A”-S1D” and Movie S3). Notably, cells photoconverted past stage 10 did not contribute to the LPM, further confirming that by this stage the contribution of the primitive streak to the LPM has ended.

**Figure 2.**
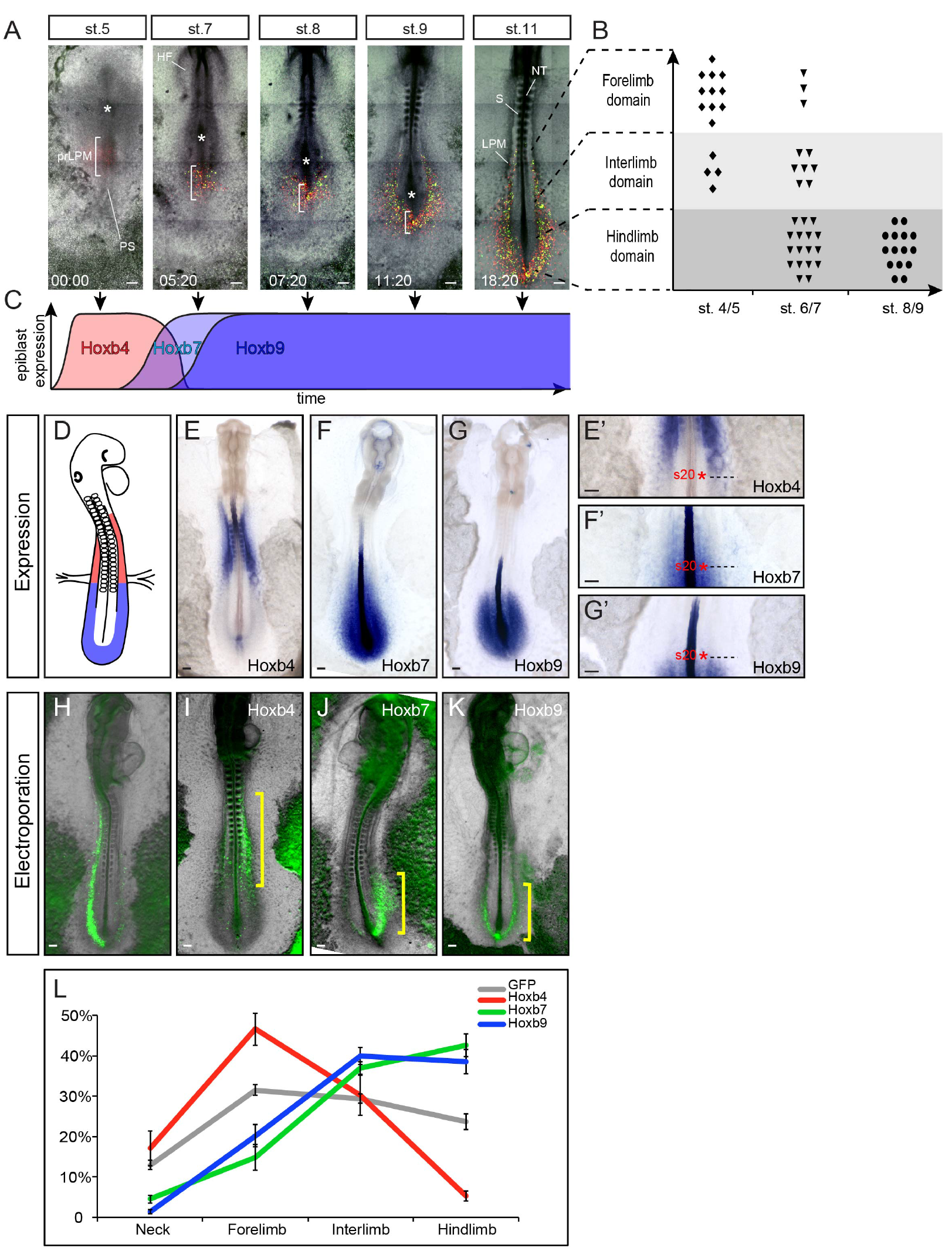
Progressive formation of the LPM and concomitant patterning by Hox genes. (A) Time series showing LPM formation from stage 5 to stage 11. LPM precursors are electroporated with H2b-RFP (red) and GFP (green). White brackets outline the presumptive LPM in the primitive streak; white asterisks, Hensen node; PS, primitive streak; prLPM, presumptive LPM; HF, head folds; NT, neural tube; S, somites. (B) Position of electroporated cells in the LPM (Y axis) as a function of their timing of ingression (X axis), n=57 tracked cells, 8 embryos. (C) Timeline of Hoxb4 (red), Hoxb7 (light blue) and Hoxb9 (blue) activation in the presumptive LPM with regard to LPM formation (black arrows). For detailed expression data see Figure S2. (D) Schematic summarizing anterior (red) and posterior (blue) Hox genes expression in the LPM. (E-G) Expression of Hoxb4 (E), Hoxb7 (F) and Hoxb9 (G) at stage 13. (E’-G’) are higher magnifications of (E-G) showing Hox genes anterior (Hoxb4, E’) or posterior (Hoxb7 and Hoxb9, F’ and G’, respectively) border of expression in the LPM (dashed black line). Red asterisks mark the prospective somite 20. (H-K) Stage 13 embryos, electroporated at stage 4 with GFP (H), Hoxb4/GFP (I), Hoxb7/GFP (J) and Hoxb9/GFP (K). Yellow brackets highlight different distribution of electroporated cells in the LPM. (L) Distribution of electroporated cells along the A-P axis for GFP-only (grey, 16 embryos, 6994 cells), Hoxb4/GFP (red, 12 embryos, 3203 cells), Hoxb7/GFP (green, 11 embryos, 2359 cells) and Hoxb9/GFP (blue, 15 embryos, 3922 cells) electroporated embryos. Error bars represent SEM. Scale bar is 100μm.

### Hox genes progressively pattern the LPM during gastrulation

During paraxial mesoderm formation, the collinear activation of Hoxb genes controls the establishment of their own expression domains through the regulation of the timing of ingression of epiblast cells [27]. Hoxb genes also display collinear activation in the prospective LPM territory of the PS, with Hoxb4 expression starting at stage 4, Hoxb7 at stage 5 and Hoxb9 at stage 6/7 (Figure S2). Importantly, activation of Hoxb4 correlates with the timing of ingression of forelimb precursor cells in the PS whereas activation of Hoxb7 and Hoxb9 genes correlates with the ingression of interlimb precursor cells (summarized in Figure 2C). By stage 11, Hox genes could be classified into 2 groups: a group of genes expressed anteriorly (anterior to the 20^th^ somite level, e.g. Hoxb4, but also Hoxb3 and Hoxb5) in a domain encompassing the prospective forelimb and another group of genes expressed posteriorly (posterior to the 20^th^ somite level, e.g. Hoxb7 and Hoxb9 but also Hoxb6 and Hoxb8) in a domain encompassing the prospective interlimb/hindlimb domain (Figures 2D-2G, 2E’-2G’and data not shown).

To test the role of different Hox genes in regulating the ingression of LPM precursor cells, we electroporated the presumptive LPM territory of the PS in stage 4 embryos either with GFP alone or in combination with Hoxb4, Hoxb7 or Hoxb9. While control GFP expressing cells were distributed uniformly along the A-P axis (Figures 2H and 2L, n=16/16), Hoxb4 expressing cells were predominantly found in the anterior part of the embryo (within the forelimb domain) (Figures 2I and 2L, n=12/14). In contrast, Hoxb7 and Hoxb9 expressing cells concentrated in the posterior-most part of the embryo (within the interlimb/hindlimb domain) (Figures 2J-2K and 2L, n=11/12 and n=15/18, respectively). Notably, cells overexpressing a given Hox gene were always found in the normal expression domain of this gene (i.e. Hoxb4 overexpression distributed anteriorly and Hoxb7/9, posteriorly; compare Figures 2I-2K, yellow brackets, with Figures 2E-2G). These results show that as for paraxial mesodermal cells [27], different Hox genes differentially regulate cell ingression of LPM precursors in the PS and, as a consequence, they control the relative position of their own expression domains in the LPM.

### Hox genes control limb initiation

These results explain how the characteristic Hox genes expression domains are positioned within the forming LPM. We next decided to test the role of these Hox-domains in instructing the forelimb position. Genes from the Hox4 and Hox5 groups have recently been proposed to control Tbx5 expression [10,11]. However, whether these Hox genes can drive endogenous or ectopic Tbx5 expression *in vivo* and whether they can modulate forelimb position has not been tested. We therefore tested whether Hoxb4, when electroporated ectopically in the interlimb region, is able to drive Tbx5 expression and displace limb position. To our surprise, as with GFP electroporation, we could not detect ectopic expression of Tbx5 in the interlimb region of embryos overexpressing Hoxb4 analyzed 24 h after electroporation (Figures 3A, 3A’ and 3B, 3B’, n= 0/20 and n=0/16, respectively). This could be due to expression of Hoxc9 in the interlimb region, as this gene can repress Tbx5 expression [11]. To test this hypothesis, we generated a truncated form of Hoxc9 lacking the C-terminal portion of the Homeodomain (required for DNA binding; for review see [28]) while conserving its repressive and paralog-specific motives, previously identified and characterized (i.e. hexapeptide motif and N terminal residues of the Homeodomain) [11] (Figure S3A). Therefore this truncated form of HOXC9 retains the ability to bind HOX9 specific and repressive co-factors but lacks the ability to bind target DNA, presumably acting as a dominant-negative (dn) form by competing for cofactors. Notably, it has been proposed that the action of such truncated HOX protein is dominant over paralogs [29–31]. In mouse, deletion of all Hox9 paralogs has been shown to prevent Sonic Hedgehog expression in the developing limb buds [32]. We validated the dominant-negative effect of the Hoxc9dn construct by electroporation in the limb mesenchyme of stage 15 chicken embryos. This led to a partial (very likely because of the mosaic efficiency of electroporation which only targets a fraction of cells), but significant decrease in Sonic Hedgehog expression, reminiscent of Hox9 mouse null mutant embryos, suggestive of a pan Hox9 inhibition (Figures S3B-S3J).

**Figure 3.**
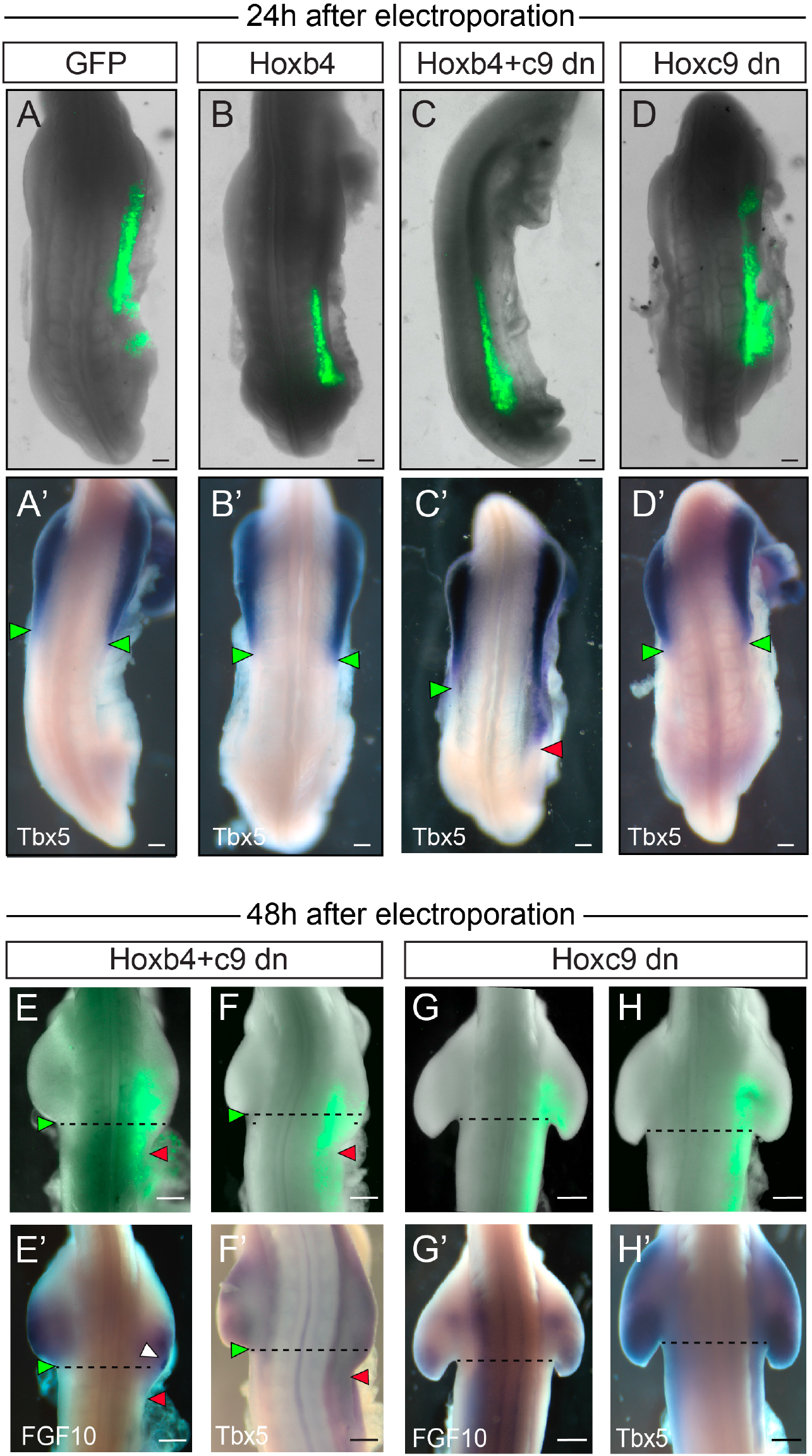
Hox genes determine limb- and non-limb forming domains within the LPM. (A-D) Embryos electroporated with GFP (A, n=20 embryos) alone or in combination with Hoxb4 (B, n=16 embryos), Hoxb4*+*Hoxc9dn (C, n=19 embryos) or Hoxc9dn (D, n=15 embryos) in the interlimb domain at stage 13 and re-incubated for 24h. (A’-D’) Tbx5 expression in corresponding embryos. Ectopic Tbx5 expression in the interlimb region is denoted by red arrowheads compare to normal endogenous expression denoted by green arrowheads (C’, n=9/19 embryos). (E-H) Embryos electroporated with GFP in combination with Hoxb4+Hoxc9dn (E, F, n=14 embryos) or Hoxc9dn (G, H, n=10 embryos) in the interlimb domain at stage 13 and re-incubated for 48h. The unilateral expansion of the forelimb is denoted by red arrowheads (E-F) compared to contralateral side denoted by green arrowhead (n=7/14 embryos). (E’-H’) shows FGF10 (E’, G’) and Tbx5 (F’, H’) expression in corresponding embryos. White arrowhead in (E’) points at ectopic FGF10 expression within the electroporated region. Scale bar is 100 m in (A-D, A’-D’) and 200’m in (E-H, E’-H’).

When Hoxb4 was electroporated in combination with Hoxc9dn and GFP, ectopic expression of Tbx5 was observed in the interlimb region (Figures 3C and 3C’, n=9/19). Notably, electroporation of Hoxc9dn alone did not promote ectopic expression of Tbx5 in the interlimb region (Figures 3D and 3D’, n=0/15), suggesting that in Hoxb4/Hoxc9dn electroporations, Tbx5 ectopic expression does not arise from the sole release of a repression imposed by the Hox9 genes. In order to observe the effects on limb position, we then allowed electroporated embryos to develop for 48h, after limb buds have clearly formed. In embryos electroporated with a combination of Hoxb4, Hoxc9dn and GFP, a posterior extension of the limb bud by 1-2 somites could be observed in 50% of the cases (Figures 3E and 3F, n=7/14). Posteriorly extended limb buds expressed the pan-limb and forelimb specific markers, FGF10 and Tbx5, respectively (Figures 3E’ and 3F’). High levels of FGF10 expression were observed in the electroporated region (Figure 3E’, white arrowhead), indicative of the ectopic activation of the limb initiation/outgrowth regulation feedback loop. As expected, neither electroporation of Hoxc9dn, Hoxb4 nor GFP alone induced a posterior extension of the limb bud (n=0/10, n=0/15 and n=0/12, respectively; Figures 3G-3H, 3G’-3H’; and data not shown). Together with previous data demonstrating Hox binding at the Tbx5 genomic locus [10,11], these results show that Hox genes regulate limb initiation through combinatorial activation and repression activities on Tbx5 expression *in vivo.*

### Relative changes in Hox collinear activation timing during gastrulation prefigure bird natural variation in limb position

Altogether these results argue that the timing of Hox activation during gastrulation defines the future positioning of their expression domain, which then determines the position of forelimb and interlimb domains. We next reasoned that natural variation in forelimb position between bird species should therefore be traced back to changes in the timing of Hox activation during gastrulation. In order to test this hypothesis, we compared the early development of three different bird species, zebra finch (Guttata taeniopygia), chicken and ostrich (Struthio camelus) whose forelimbs begin at the level of the 1^st^ thoracic vertebrae which is the 13th, 15th and 18th vertebrae, respectively (Figures 4A-4C). Surprisingly, when compared to chicken embryo, the Tbx5-positive forelimb field in embryos of these species did not reveal a translation of the Tbx5 expression domain (Figure 4D-4F) but instead an anterior (of about 3 somites) and a posterior (of about 5 somites) shift of its posterior border only, in zebra finch (Figures 4E, 4E’) and ostrich embryos (Figures 4F, 4F’), respectively. As a result, the forelimb field is gradually extended when comparing embryos of these species. A concomitant shift of the Hoxb4/Hoxb9 border by about 3-5 somites could be observed anteriorly in zebra finch (Figures 4H-4H’ and 4K-4K’) and posteriorly in ostrich embryos (Figures 4I-4I’ and 4L-4L’), when compared to chicken embryos (Figures 4G and 4J). Therefore, a shift in the Hoxb4/Hoxb9 border and in the posterior border of Tbx5 foreshadows the differences in limb position.

**Figure 4.**
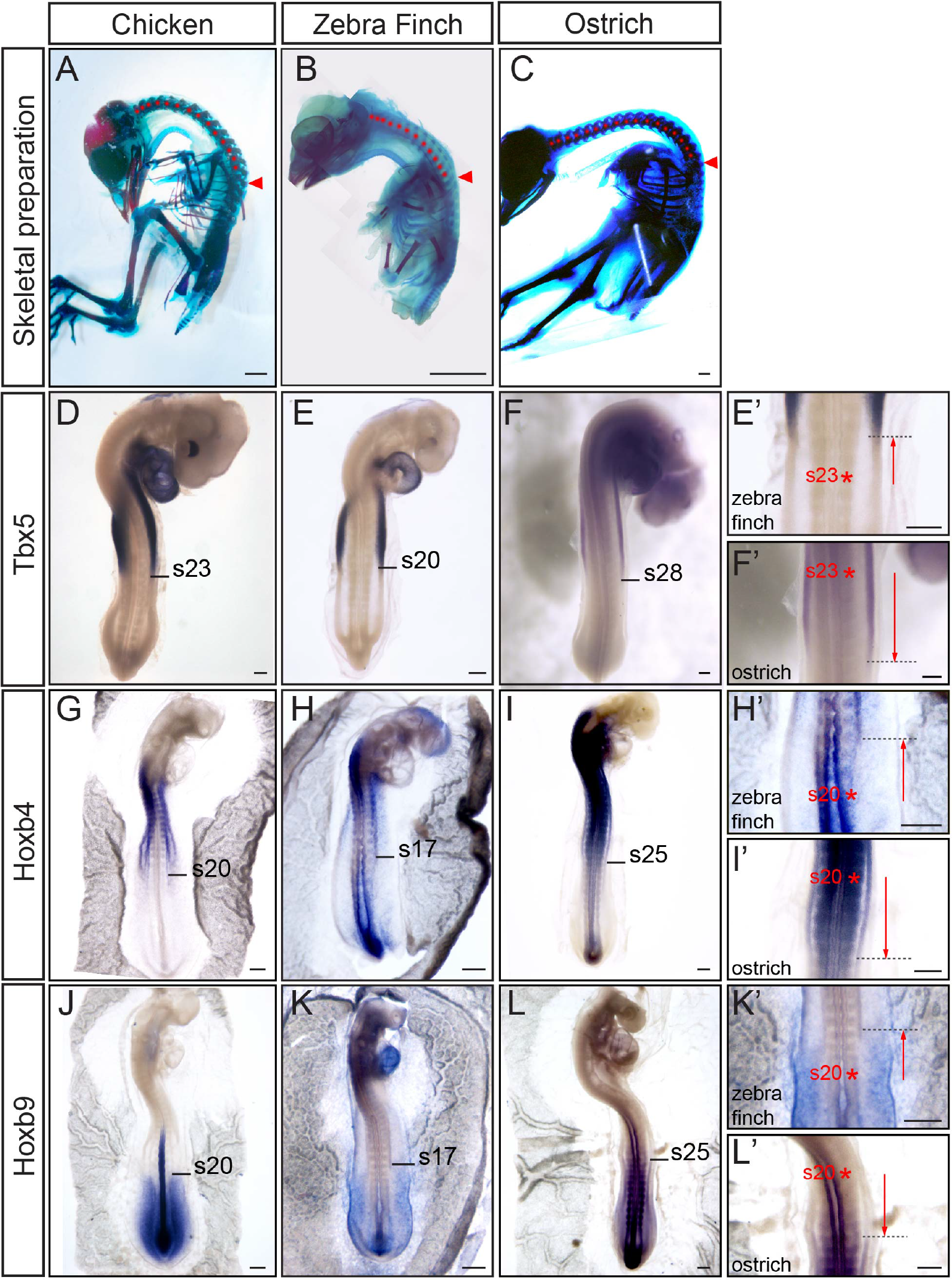
Variations in wing position correlate with changes in Tbx5, Hoxb4 and Hoxb9 expression domains in the LPM. (A-C) Alcian Blue/Alizarin Red staining of chicken (A, E20), zebra finch (B, E13) and ostrich (C, E37). Red arrowheads point at the wing position level (15^th^, 13^th^ and 18 ^th^ vertebrae in chicken, zebra finch and ostrich embryos, respectively); red dots mark each cervical vertebrae. (D-F) Tbx5 expression in stage 18 chicken (D) zebra finch (E) and ostrich embryos (F). The position of Tbx5 posterior border of expression is indicated in somite number. (E’) and (F’) are higher magnification of (E) and (F), respectively. (G-I) Hoxb4 expression in chicken (G, 20-somite stage), zebra finch (H, 20-somite stage) and ostrich embryos (I, 34-somite stage). The position of Hoxb4 posterior border of expression is indicated in somite number. (H’) and (I’) are higher magnification of (H) and (I), respectively. (J-L) Hoxb9 expression in chicken (J, 20-somite stage), zebra finch (K, 20-somite stage) and ostrich embryos (L, 36-somite stage). The position of Hoxb9 anterior border of expression is indicated in somite number. (K’) and (L’) are higher magnification of (K) and (L), respectively. Dashed black lines show variation in posterior/anterior border of expression in zebra finch (E’, H’ and K’) and ostrich embryos (F’, I’ and L’) compared to chicken embryo (represented by red asterisks). Scale bar is 3mm in (A-C) and 100mm (D-L, E’-F’, H’-I’ and K’-L’).

We then investigated whether temporal differences in collinear activation of Hoxb4 and Hoxb9 genes during gastrulation could be linked to the spatial variation of the Hoxb4/Hoxb9 border in chicken and ostrich embryos. We found that Hoxb4 is activated at the same stage (stage 4) in both chicken and ostrich embryos (Figures 5A and 5E). However, Hoxb4 remained expressed in the epiblast for much in longer in ostrich (10-somite stage) than in chicken embryos (2-somite stage) (Figures 5B, 5F and S2C). Concomitant to this delay in Hoxb4 arrest of expression in the epiblast, Hoxb9 activation was delayed in ostrich (10-somite stage, Figures 5N-5P) compared to chicken embryos (2-somite stage, Figures 5J-5K). Ultimately the Hoxb4/Hoxb9 border became posteriorly shifted in ostrich embryos when compared to chicken embryos, although at a later stage (Figures 5C-5D, 5G-5I, 5L-5M and 5Q-5R). Altogether these results show that relative changes in the timing of Hoxb genes activation and in particular the transition between anterior (Hoxb4) and posterior (Hoxb9) Hox genes prefigure spatial variation of Hox genes expression domains and limb position observed at later stages (summarized in Figure 5S).

**Figure 5.**
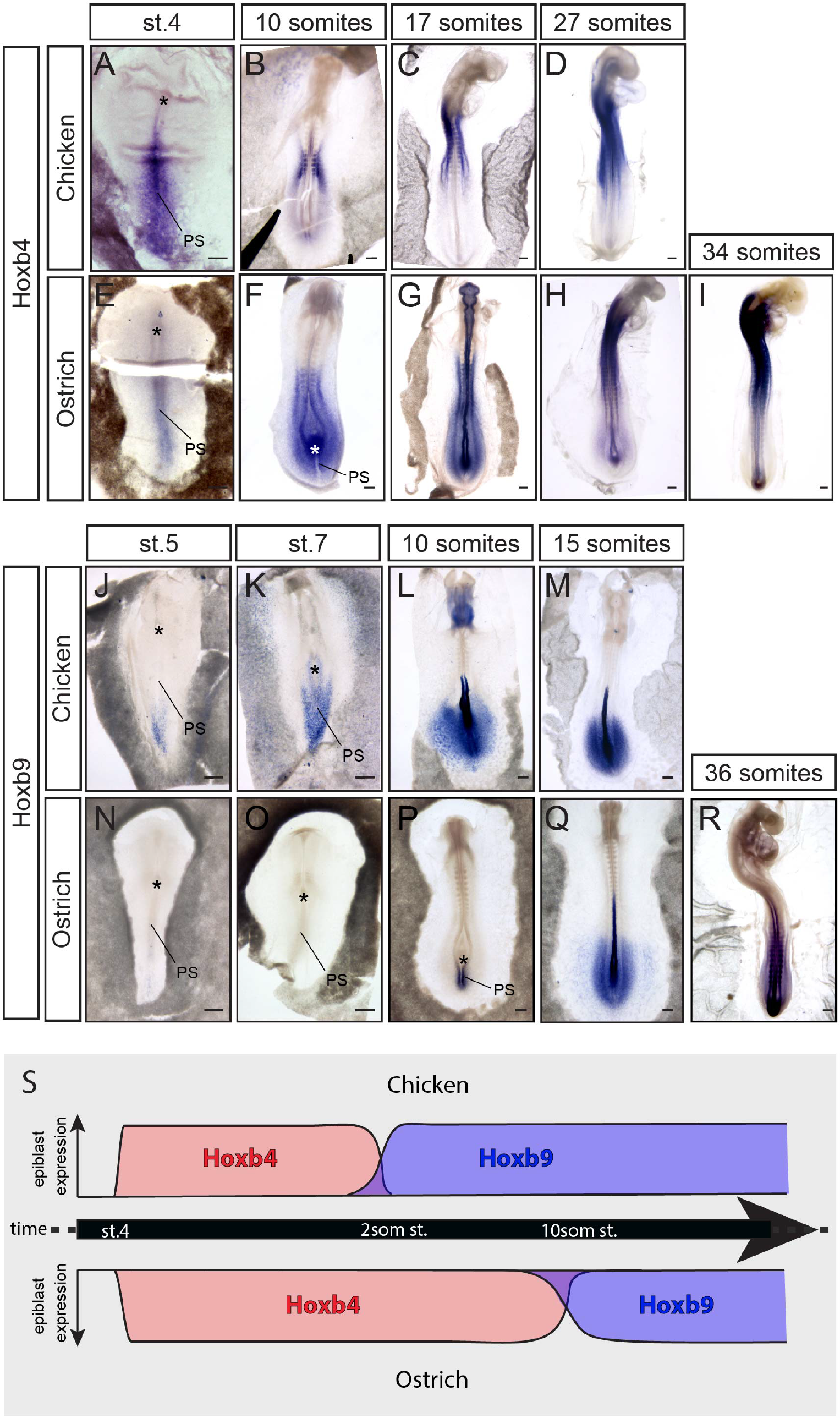
Relative changes in Hox collinear activation timing underlie variation in limb position between chicken and ostrich. (A-I) Hoxb4 expression in chicken (A-D) and ostrich (E-I) embryos. (J-R) Hoxb9 expression in chicken (J-M) and ostrich (N-R) embryos. (S) Timeline of Hoxb4 (red) and Hoxb9 (blue) activation in chicken (top diagram) and ostrich (bottom diagram). Scale bar is 100μm. Asterisks represent the Hensen node; PS, primitive streak.

### Retinoic Acid signaling modulation during gastrulation changes the extent of the forelimb field

We next sought to understand how variation in Hox activation timing is controlled between zebra finch, chicken and ostrich. Differences in the timing of expression of Gdf11 have recently been proposed to account for variation in hindlimb position between tetrapods [33]. Since Gdf11 acts through modulation of Retinoic Acid (RA) signaling by inducing Cyp26a1 expression (a RA catabolizing enzyme) [34] and because RA was shown to activate anterior Hox gene in the neural tube [35] and repress posterior Hox genes in the tail bud [36], Cyp26a1 stood as an excellent candidate to regulate variation in Hox activation timing. We therefore compared Cyp26a1 expression between the different species and found that it is first detected in the prospective LPM territory of the PS at stage 4 in zebra finch, stage 5-6 in chicken and stage 10 in ostrich (Figures 6A-6C, red arrowheads). Therefore Cyp26a1 is precociously expressed in zebra finch and delayed in ostrich compared to chicken embryo. Notably, the onset of Cyp26a1 expression in the primitive streak precedes by a few hours the transition between Hoxb4 and Hoxb9 expression in the epiblast.

**Figure 6.**
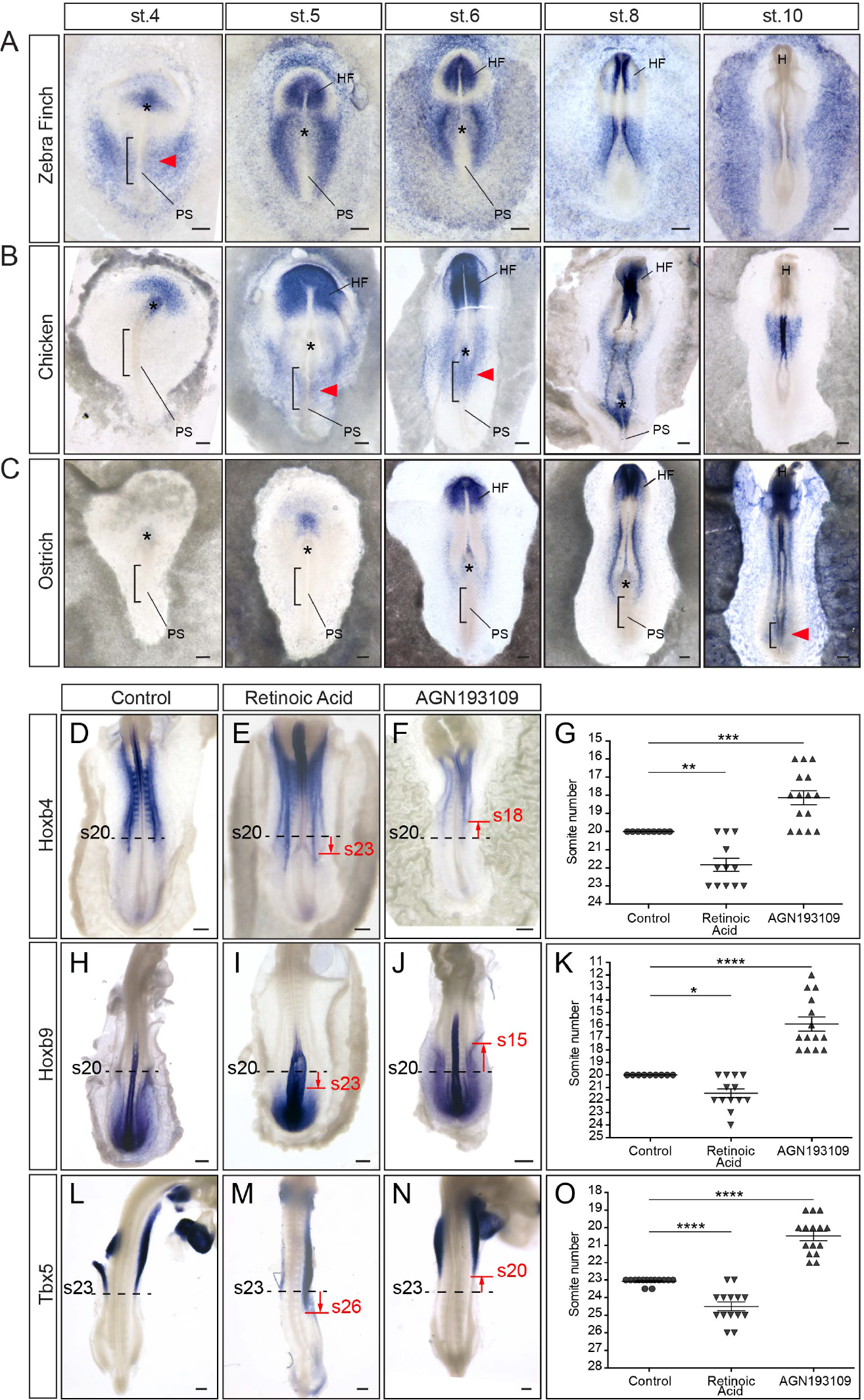
Changes in Retinoic Acid signaling during gastrulation modulate forelimb field position. (A-C) Cyp26a1 expression in zebra finch (A), chicken (B) and ostrich (C) embryos. Red arrowheads highlight the onset of Cyp26a1 expression in the presumptive LPM (black brackets); black asterisks, Hensen node; PS, primitive streak; HF, head folds; H, head. (D-F) Hoxb4 expression in stage 13 control (D), Retinoic Acid-(E) and AGN193109- (F) treated embryos. (G) Position of Hoxb4 posterior border of expression in control (n=9 embryos), Retinoic Acid- (n=12 embryos) and AGN193109-treated embryos (n=15 embryos). (H-J) Hoxb9 expression in stage 13 control (H), Retinoic Acid- (I) and AGN193109- (J) treated embryos. (K) Position of Hoxb9 anterior border of expression in control (n=9 embryos), Retinoic Acid- (n=13 embryos) and AGN193109-treated embryos (n=14 embryos). (L-N) Tbx5 expression in stage 18 control (L), Retinoic Acid- (M) and AGN193109- (N) treated embryos. (O) Position of Tbx5 posterior border of expression in control (n=14 embryos), Retinoic Acid- (n=14 embryos) and AGN193109-treated embryos (n=15 embryos). Red lines show changes in posterior/anterior border of expression in treated embryos compared to control embryos (dashed black lines). ANOVA with Fisher LSD post hoc test, * p<0,05; ** p<0,01; *** p<0,001; **** p<0,0001. Scale bar is 100μm.

Finally, we tested whether modulating Retinoic Acid signaling during gastrulation could alter the spatial organization of Hox expression domains and the forelimb field position. When stage 4 chicken embryos were incubated in presence of RA or AGN193109, a pan Retinoic Acid Receptor antagonist, the posterior border of Hoxb4 became shifted of about 2-3 somites posteriorly and anteriorly, respectively (Figures 6D-6G). A complementary posterior and anterior shit of Hoxb9 anterior border was observed in embryos treated with RA and AGN193109, respectively (Figures 6H-6K). Therefore the modulation of RA signaling during gastrulation changes the relative position of the Hoxb4/Hoxb9 border in the LPM, at later stages. We next checked the effect on Tbx5 expression and observed a concomitant shift of its posterior border of about 2-3 somites posteriorly in embryos treated with RA and anteriorly when these embryos were treated with AGN193109 (Figures 6L-6O). Notably, this led to changes in the extent of Tbx5-positive forelimb field in RA and AGN193109-treated embryos which strikingly resemble the extent of forelimb fields in ostrich and zebra finch embryos, respectively (compare Figure 6M and 6N with Figure 4F and 4E, respectively). Altogether these results show that modulation of Retinoic Acid signaling during gastrulation affects the axial extent of Hox expression domains in the lateral plate mesoderm and provokes modification in the A-P extent of Tbx5 expression, which is linked to variation in limb position.

## Discussion

Our work identifies an early role for Hox genes in the regulation and variation of forelimb position in birds. During gastrulation, the lateral plate compartment is progressively generated and concomitantly patterned by Hox genes. As observed for the adjacent paraxial mesoderm, collinear activation of Hox genes themselves in the epiblast progressively establishes their own collinear domains of expression in the LPM. It is the position of these expression domains that determines the position of forelimb-forming and interlimb-forming domains through a combination of activation (e.g. Hoxb4) and repression (e.g. Hox9) of limb initiation (i.e. Tbx5 expression). Furthermore, relative changes in the collinear activation of Hox genes during gastrulation prefigure variation in the spatial organization of Hox genes expression domains and natural variation in limb position between birds. We also observe differences in the onset of expression of the RA catabolizing enzyme Cyp26a1 during gastrulation and show that modulation of RA signaling provokes changes in limb field position. Based on our findings we therefore propose that timely controlled Hox collinear activation during gastrulation is responsible for the regulation and variation in forelimb position in birds.

### Progressive formation and patterning of the LPM by collinear activation of Hox genes

Our results show that the LPM compartment is progressively formed during gastrulation. Surprisingly, despite a thorough characterization of other embryonic derivatives adjacent to the prospective LPM in the epiblast, the cellular events underlying the formation of the LPM during gastrulation, to the best of our knowledge, had never been described. Whereas, the prospective territory of the LPM had been traced back to the middle third of the primitive streak [15,16,19], how forelimb, interlimb and hindlimb arise from this field was not addressed. Here we show that these fields sequentially form between stage 4 and 10, (between 24h and 48h of development). More importantly, by characterizing the precise timing of forelimb, interlimb and hindlimb domain formation, we could link the formation of these domains to the concomitant collinear Hox gene activation. By comparing Hox activation in zebra finch, chicken and ostrich embryos, we further show that relative differences in the timing of Hox activation prefigure spatial variation of their domain of expression as predicted by experiments performed in chicken. Previous studies have shown that in the context of paraxial mesoderm formation, Hox genes, collinearly activated, establish through the regulation of cell ingression at the streak, their own spatial collinear characteristic pattern [27]. We show that a similar mechanism is at work during the generation of LPM and further show the implication of such process on subsequent patterning (limb- and non-limb-forming domains) but also on variation in body plan organization. The finding that paraxial and LPM are similarly generated and patterned during gastrulation provides a simple explanation for the concomitant patterning of the cervico-thoracic frontier in the somites and the associated forelimb position in LPM.

In birds, the variation of limb position is produced by meristic variations (characterized by changes in the total number of component parts; [37]) meaning that this variation is accompanied with the addition of segmental units (somites) along with LPM tissue. As an example, whereas sparrows have 9 cervical vertebrae, swans exhibit 16 additional vertebrae. These meristic variations imply that at the embryonic level a larger amount of mesodermal cells have to be produced to form both the somitic and corresponding LPM tissue. A recent study has linked the collinear activation of posterior Hox genes (Hox9 to 13) to the regulation of axis elongation and its termination. The temporal collinear activation of posterior Hox genes was shown to slow down the influx of mesodermal cells through the primitive streak in a collinear trend thereby controlling the elongation rate [29]. Coupling both the patterning of mesodermal tissues and the rate of axis elongation to the same regulation mechanism might therefore ensure that addition of somites and corresponding LPM tissue is not made at the expense of the subsequently formed somites, therefore maintaining the vertebrate body integrity and providing a simple mechanism to control meristic variations observed in birds.

### Hox gene expression determine limb and non-limb forming domains

The role of Hox genes in the determination of limb position has been put into question, given the lack of clear phenotypes in Hox genes loss- and gain-of-function experiments in mouse [38]. More recently, evidences reinforcing the role of Hox genes in positioning limb fields have accumulated. Hox genes from paralog groups 4 and 5 can bind to a Tbx5- forelimb specific enhancer and activate transcription of a downstream reporter [10,11], however this has not been shown *in vivo*, on the endogenous Tbx5 expression. Intriguingly, whereas Tbx5 expression is greatly conserved among vertebrates, the above-mentioned Tbx5 limb-specific enhancer, located in intron 2 in mouse, could not be located in chicken, raising doubts about the requirement of this particular enhancer for Tbx5 forelimb expression. Moreover, the fact that Tbx5 has recently been shown to be insufficient (although necessary) for forelimb initiation [39] also raised concerns about the sufficiency of displacing forelimb position by solely displacing Tbx5 domain. Hoxc9 in turn, was shown to bind directly Tbx5 limb enhancer and to inhibit its expression, as a consequence it has been proposed that there is a latent potential in the caudal LPM to express Tbx5 that is normally masked by the presence of Hoxc8-10 genes [11,40]. Our data which shows that ectopic expression of Hoxb4, in combination with a Hoxc9 dominant negative, can induce Tbx5 expression and extend the forelimb is in full support with a role of Hox4-5 genes in regulating Tbx5 expression *in vivo*. However, the finding that Hoxc9dn on its own does not promote Tbx5 expression nor forelimb expansion but does it only in combination with Hoxb4, has several important implications with respect to regulation and variation of limb position. First, it suggests that there is no latent potential of forelimb forming activity in the interlimb but “just” a repressive forelimb-forming activity. Second and corollary to this, it suggests that a posterior shift in limb position must not only involve a shift of the forelimb field (e.g. Hoxb4 expression, which is normally not expressed posteriorly) but also of the interlimb field (e.g. Hoxb9 anterior border of expression). In other words, changes in limb position can be induced only if the overall spatial sequence of Hox expression pattern is changed. This might explain why none of the single nor compound mutants for a variety of Hox genes including Hox5 and Hox9 groups show a major phenotype on limb position [32,41]. Indeed, our data argues that to induce a shift in forelimb position in mouse, a combination of gain- and loss- of function for forelimb-activator and forelimb-repressor Hox genes, respectively, should be performed. Based on these results, we propose that the sequence limb-forming/non-limb-forming domains is embedded in the collinear organization of Hox genes through their specific activation/repression function on limb initiation, it is in turn the timing of collinear activation that sets the relative position of these domains along the main axis.

### Posterior border of the early limb field and final limb position

Our results point at the posterior border of the forelimb (Hoxb4/Hoxb9 border) as a critical regulator of limb positioning since only the position of the posterior (and not the anterior) border of Hoxb4 and Tbx5 is gradually shifted in zebra finch, chicken and ostrich. This automatically leads to a gradual extension of the forelimb field rather than a gradual posterior translation, although the limb becomes eventually posteriorly shifted between these species. How can a variation in forelimb field size lead to variation in limb position? Interestingly, it has been shown that Tbx3 can alter the position of the forelimb through an interplay with Gli3 and dHand gradients which prepattern the early mesenchyme and set the position of the ZPA [42]. Modulation of this prepatterning (by modulating Tbx3 activity) subsequently modifies the position of the ZPA and eventually the position of the limb [42]. Strikingly, Hox5 mutants in mice show derepression of Shh, expressed in the ZPA [41], whereas in Hox9 mutant, Shh is not expressed due to defects in Gli3/dHand prepatterning of the early mesenchyme [32]. It is thus tempting to speculate that once Hox genes have set the forelimb field of variable size in different species, prepatterning of the early mesenchyme by Hox5, Hox9 and most likely other Hox genes could in turn induce variation in the position of the ZPA which could be responsible for variation in the definitive position of the forelimb. Notably, the fact that Tbx5 activation upon electroporation of Hoxb4 and Hoxc9dn does not induce an ectopic limb bud but rather its expansion is in agreement with concomitant re-patterning of this larger field and also supports recent report of Tbx5 insufficiency in promoting (ectopic) limb bud formation [39] as opposed to what was previously reported [43].

### Retinoic Acid signaling as regulator of both forelimb and hindlimb position?

We show that Cyp26a1 expression during gastrulation is preciously expressed in zebra finch and delayed in ostrich when compared to chicken and correlates with the activation of interlimb specific Hox genes such as Hoxb7 or Hoxb9 in each of these species. Furthermore, we find that modulation of Retinoic Acid signaling during gastrulation leads to changes in the extent of Hox gene expression patterns while preserving overall collinearity. Consequently, we observe a variation in the posterior border of the forelimb domain, as revealed by Tbx5 expression, which recapitulates natural variation observed between zebra finch, chicken and ostrich. The possibility that Cyp26a1, which has been implicated in positioning hindlimbs [34], might regulate forelimb position is particularly interesting since a single signaling pathway would therefore be responsible for the regulation of both forelimb and hindlimb position, although at different developmental timing.

## Acknowledgements

We thank Francois Schweisguth and Cliff Tabin for critical reading of the manuscript; Marie Manceau for providing zebra finch fertilized eggs. CM was supported by a MENRT fellowship. The research leading to these results has received funding from the European Research Council under the European Union’s Seventh Framework Programme (FP7/2007-2013) / ERC Grant Agreement *n°337635, from the* Institut Pasteur, the CNRS and the Cercle FSER.

## Author Contribution

Conceptualization, CM and JG; Methodology, DR and JR; Investigation, CM, PC, JG; Writing – Original Draft, JG; Writing – Review and Editing, JG, CM, OP; Resources, ND and OP; Funding acquisition, JG; Supervision, JG.

## Material and Methods

### Avian Embryos

Fertilized chicken and ostrich eggs were ordered from commercial sources (chicken: EARL Morizeau; ostrich: SARL Le père Louis), fertilized zebra finch eggs were generously provided by Dr Marie Manceau from Collège de France (Paris) and transgenic quail eggs were produced in the lab. Chicken, quail and zebra finch eggs were incubated at 38°C and ostrich eggs at 36°C. All embryos were staged according to the Hamburger and Hamilton classification system [44].

### Microsurgery

Microsurgery experiments were performed in stage 11 embryos. The ectoderm was carefully detached in order to only microdissect the right somatopleure encompassing both forelimb and interlimb prospective domains, as previously proposed [20,45]. This tissue was rotated along its A-P axis and grafted back into the same embryo. Eggs were further sealed with tape and reincubated at 38°C. After 48h, embryos were harvested and fixed in 4% formaldehyde. A similar procedure was used to perform quail-chick chimeric grafts. The stage 11 *hUbC:memGFP* transgenic quail embryo was collected and placed into a petri dish filled with PBS, the somatopleure was microdissected as described above and grafted to a stage 11 host chicken embryo.

### Embryo Culture, *ex ovo* Electroporation and Imaging

#### Embryo culture

Embryos were prepared for *ex ovo* culture using a modified version of the EC culture system [26]. Embryos were collected at stage 4 and placed in a Petri dish with semisolid nutritive medium. Embryos were then incubated at 38°C in a humidified chamber and cultured for up to 48h.

#### ex ovo Electroporation

Embryos were electroporated in a custom-made electroporation chamber using the SuperElectroporator NEPA21 type II® (NEPAGENE) with two 5ms poring pulses of 15V, 50ms delay, and three 50ms transfer pulses of 10V, 500ms delay. Solutions of plasmid DNA were prepared as previously described [46] with a final DNA concentration of 1μg/μl or 5μg/μl.

#### Imaging

Stage 4 embryos co-electroporated with pCAGGS-H2b-RFP (1μg/μl), pFlox-pA-EGFP (1μg/μl) and pCX-Cre (50ng/μl) plasmid DNAs, were placed into medium-containing glass-bottom Petri dishes (MatTek). Embryos were then imaged at 38°C using an inverted two-photon microscope (Zeiss, NLO LSM 7MP) coupled to a Chameleon Ti/Saph laser and OPO system (Coherent) at 840nm (to image GFP) and at 1100nm (to image RFP) with 10x long-distance objectives. Embryos were acquired every 5min for about 24 hours using the tiling/stitching feature of the Zen software (Zeiss).

Photoconversion of the *hUbC:mEOS2FP* transgenic quail embryos was performed using an inverted confocal microscope (ZEISS LSM 880). Before photoconversion, mEOS2 was visualized under standard imaging conditions for GFP using a 488nm Argon laser (15% power). A 405nm diode laser (40% power with 1×25 iterations) was used for photocoversion. Photoconverted mEOS2 was subsequently imaged using a 561nm HeNe laser (30% power). Embryos were imaged every 2 hours to follow the photoconverted region of the embryo, without bleaching.

### *In ovo* Electroporation and Drug administration

*In ovo* electroporation was performed as previously [46]. Briefly, eggs were incubated up to stage 13 (50h incubation). DNA solution was injected in the coelomic cavity at the level of the forelimb and interlimb domains. Electroporation was performed using homemade electrodes and the SuperElectroporator NEPA21 type II® (NEPAGENE) with two 1ms poring pulses of 70V, 100ms delay and three 2ms transfer pulses of 40V, 500ms delay. Eggs were sealed with tape and reincubated for 24h or 48h. Embryos were then harvested and fixed in 4% formaldehyde.

Chicken embryos treated with Retinoic Acid (Sigma, 500μM in DMSO) and AGN193109 (Tocris, 100μM in DMSO), were incubated up to stage 4 and drugs were injected in between the vitelline membrane and the embryo, on top of the primitive streak. Note that drug concentration below this level did not provide effects. Control embryos were treated with equivalent concentration of DMSO. Eggs were sealed with tape, reincubated 24h or 48h, embryos were then collected at stage 13 (19-22somites) and stage 18 (30-36 somites), respectively and fixed in 4% formaldehyde.

### DNA constructs

Hoxb4, Hoxb7 and Hoxb9 were subcloned into the pCAGGS-IRES2-Venus vector [27,29] and were used alone or in combination with pCAGGS-GFP. The constructs pFlox-pA-EGFP and pCX-Cre were kindly provided by X. Morin [47] and used in combination with pCAGGS-H2b-RFP. The truncated form of HOXC9 was generated by inserting a stop codon instead of the highly conserved tryptophan amino acid on the α-helix III of the Homeodomain as previously described [29–31] and further subcloned into the pCAGGS-IRES2-Venus vector.

### Immunofluorescence

Embryos were embedded in 7.5% gelatin/15% sucrose and sectioned using a Leica CM3050S cryostat. Sections were then incubated with Alexa Fluor™ 555 Phalloidin (1:100, Invitrogen) and rabbit anti-GFP primary antibody (1:500, Torrey Pines Biolabs) overnight, washed for 24h, incubated 2h with goat anti-rabbit Alexa Fluor™ 488 secondary antibody (1:1000, Invitrogen) and washed for 24h. All incubations and washes were performed in PBS/BSA 0.2%, Triton 0.1%/SDS 0.02%. Sections were then mounted with DAPI-containing Fluoromount-G™ and imaged using an inverted confocal microscope (Zeiss LSM700).

### Image analysis

#### Cell tracking

Time-lapse movies of embryos electroporated with cytoplasmic GFP and nuclear H2b-RFP fluorescent reporters were analyzed using ImageJ software and the retrospective cell tracking was performed using the Manual Tracking plugin from Fabrice Cordelières.

#### Electroporated cell distribution in the LPM

Embryos were electroporated at stage 4 with GFP, GFP/Hoxb4, GFP/Hoxb7 or GFP/Hoxb9, cultured *ex ovo* and harvested at stage 13. The LPM was segmented along the A-P axis into neck, forelimb, interlimb and hindlimb regions. Using ITCN (Imaged-based Tool for Counting Nuclei) plugin from Thomas Kuo and Jiyun Byun (UC Santa Barbara) in ImageJ software, the number of electroporated cells was quantified in each region and normalized relative to the area of these regions.

### *In situ* hybridization

*In situ* hybridization on chick embryos was performed as previously described [48]. DIG-labeled probes were amplified from plasmids containing cDNA fragments of cHoxb4, cHoxb7, cHoxb9 (as previously published [27]), cTbx5, cFgf10, cCyp26a1 and cShh.

### Skeleton analysis

Chicken, zebra finch and ostrich embryos were collected at late development stage (chicken E20; ostrich E37; zebra finch E13) and stained with Alcian Blue and Alizarin Red to label cartilage and bone tissues, respectively, as previously described [49].

### Generation of transgenic quail lines

Two transgenic lines were created in this study (*hUbC:memGFP* and *hUbC:mEOS2FP*) by following previously published method [50]. Briefly, non-incubated quail eggs (*Coturnix japonica*) were windowed and a solution of high titer lentivirus was injected into the subgerminal cavity of stage X embryos. Eggs were sealed with a plastic piece and paraffin wax. Injected eggs were incubated at 37.5°, 56% humidity until hatching. For the *hUbC:memGFP* line, a total of 42 embryos were injected with the lentivirus solution (titer 10^10^/ml). Three F0 mosaic founder males successfully hatched and reached sexual maturity (7%). They were bred to WT female and all three produced transgenic offspring (transmission rate: 8.8%). One line was selected on the basis of a single copy of the transgene, checked by Southern Blot, and high intensity of the memGFP signal. For the *hUbC:mEOS2FP* line, a total of 141 embryos were injected with lentivirus (titer 6,4.10^10^/ml). Five F0 mosaic founder successfully hatched and reached sexual maturity (3.5%). All five produced transgenic offspring (transmission rate: 6.1%) and one line was selected by Southern Blot analysis for single transgene integration and high intensity of the mEOS2 fluorescent signal.

## Supplemental Information

### Supplemental Figures

**Figure S1.**
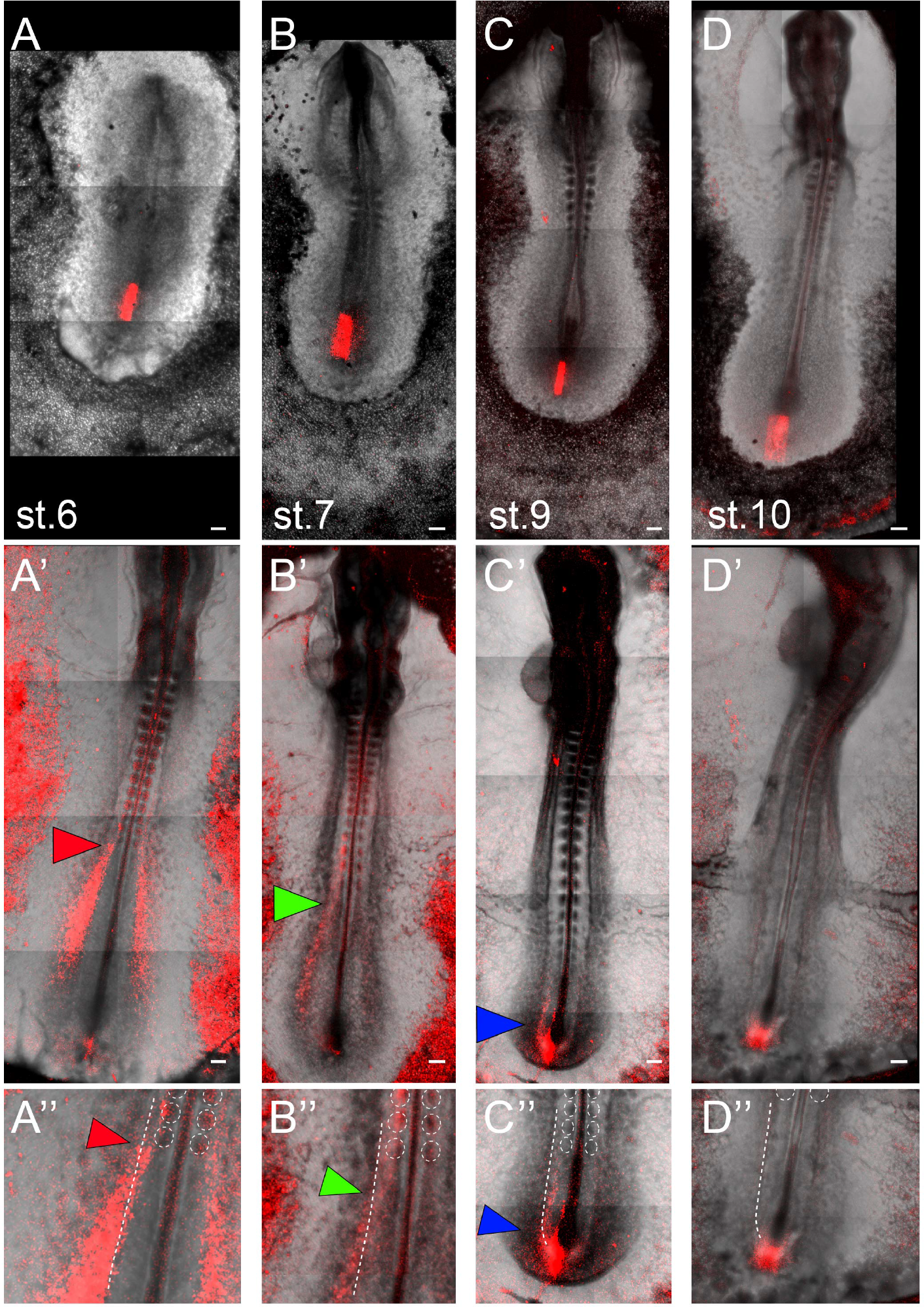
Related to Figure 2. Progressive formation of the LPM during gastrulation. (A-D) Photoconversion (red) of the prospective LPM in the primitive streak of *hUbC:mEOS2FP* transgenic quail embryos at stage 6 (A), 7 (B), 9 (C) and 10 (D). (A’-D’) Distribution of red-labeled cells 24h after photoconversion. Cells photoconverted at stage 6 (A) are found in the forelimb and interlimb domains (A’, red arrow head). Cells photoconverted at stage 7 (B) are found more posteriorly, in the interlimb and hindlimb domains (B’, green arrowhead). Cells photoconverted at stage 9 (C) are only found in the hindlimb domain (C’, blue arrowhead). Photoconversion of the remaining primitive streak at stage 10 (D) only labels the tail bud tissue (D’). Note the gradual posterior distribution as cells are photoconverted at later stages. (A”- D”) Higher magnification of (A’-D’) showing the anterior limit of photoconverted cells (arrowheads). Dashed white lines outline the border between lateral plate and somitic mesoderm; dashed white circles outline somites. n=16 photoconverted embryos. Scale bar is 100μm.

**Figure S2.**
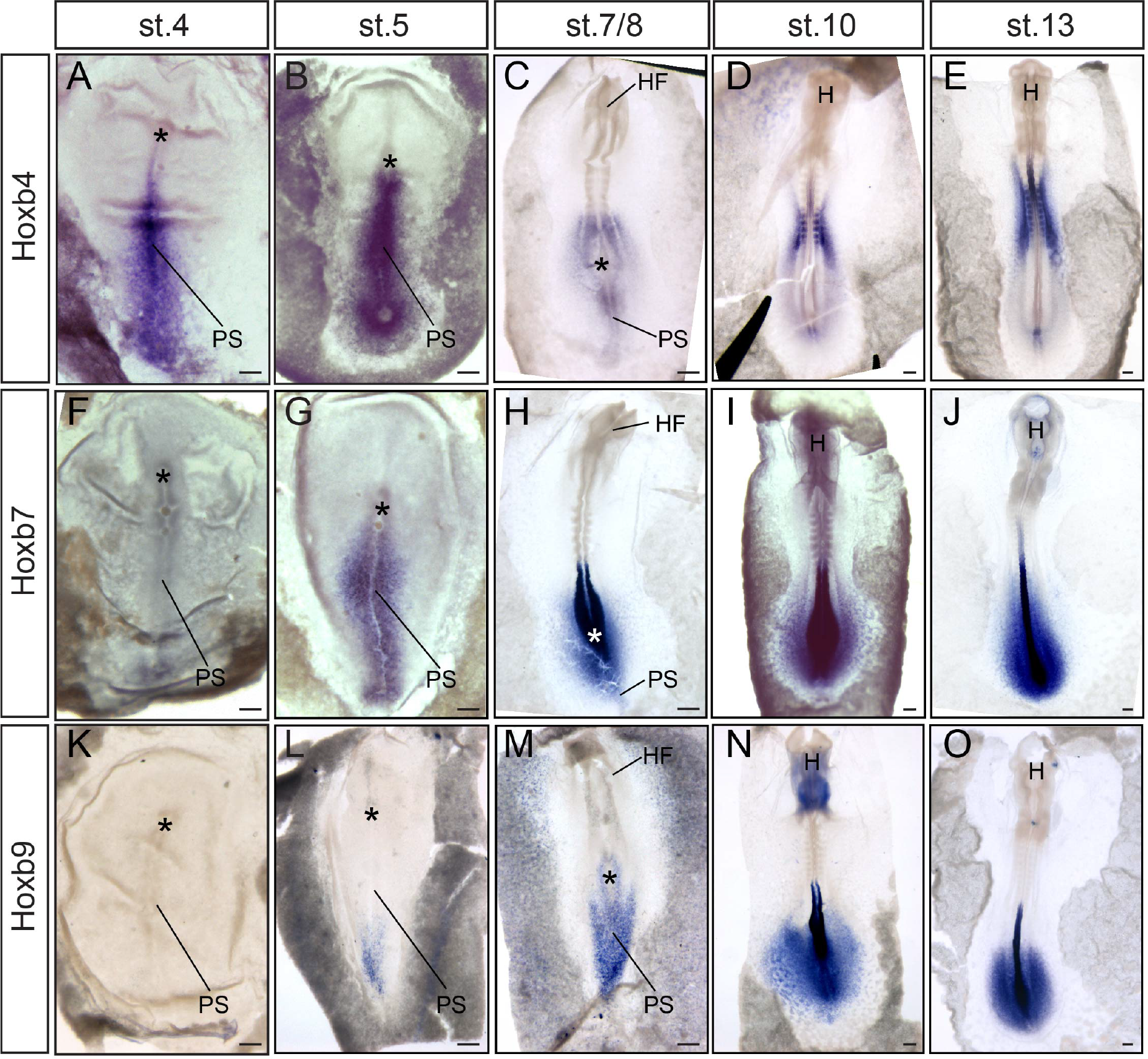
Related to Figure 2. Characterization of Hoxb temporal and spatial collinearity with respect to the LPM. (A-O) Hoxb4 (A-E), Hoxb7 (F-J) and Hoxb9 (K-O) expression patterns. Asterisks represent the Hensen node; PS, primitive streak; HF, head folds; H, head. Scale bar is 100μm.

**Figure S3.**
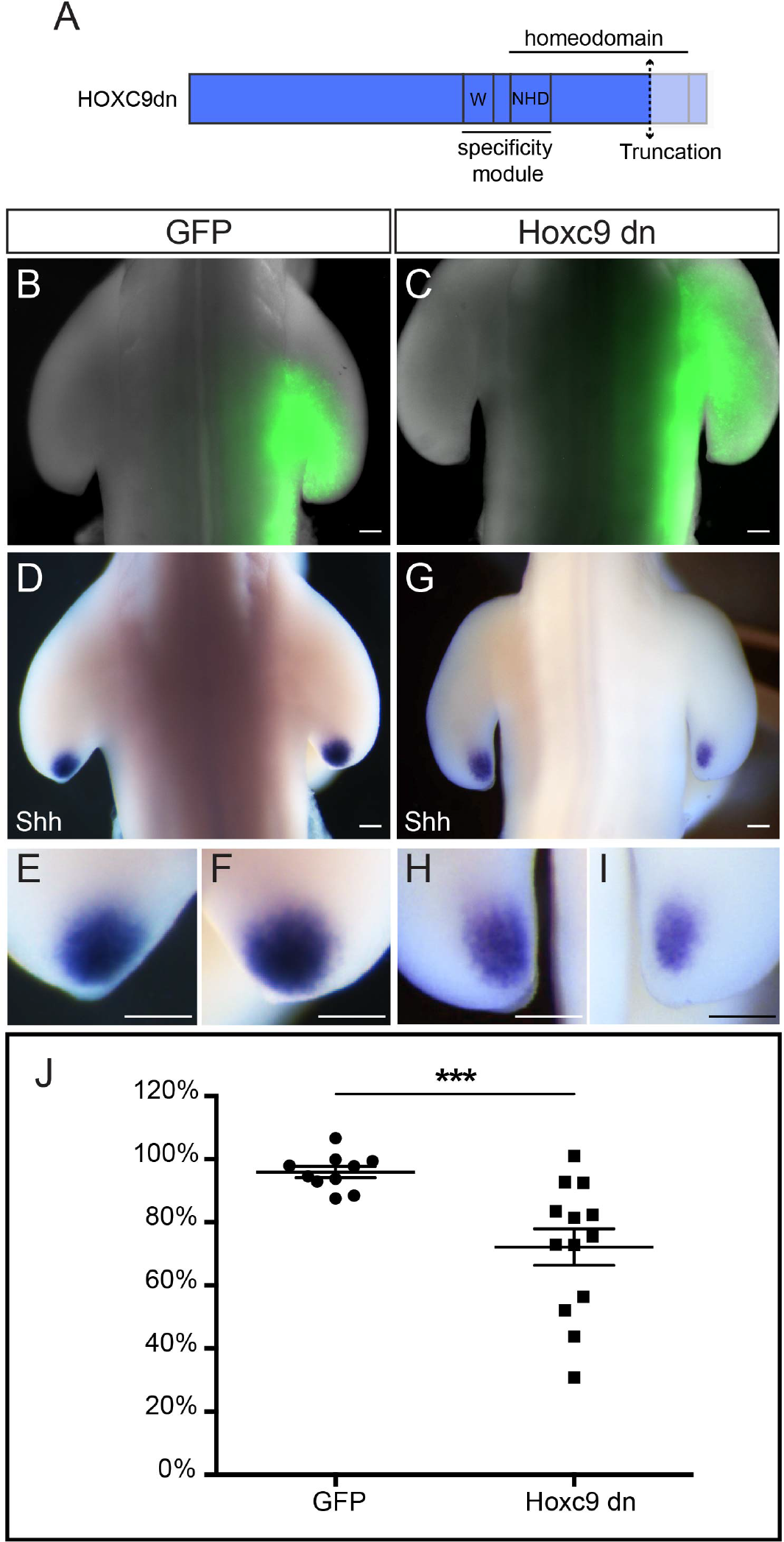
Related to Figure 3. Hoxc9dn downregulates Sonic Hedgehog (Shh) expression in the posterior limb mesenchyme. (A) Schematic representation of the truncated HOXC9 (HOXC9dn) protein. The truncated C-terminal part of the protein (including the C-terminal part of the Homeodomain) is highlighted by the dashed double arrow and the light blue shaded area. The specificity module includes the hexapeptide motif (W) ant N-terminal residues of the Homeodomain (NHD) as previously characterized [11]. (B-C) Embryos electroporated with GFP alone (B, n=10 embryos) or in combination with Hoxc9dn (C, n=13 embryos) in the forelimb domain at stage 15 and re-incubated for 48h. (D-F) Shh expression (D) in a GFP electroporated embryo (B). (E) and (F) are higher magnifications of the posterior limb mesenchyme on the contralateral (E) and GFP electroporated (F) sides. (G-I) Shh expression (G) in a GFP/Hoxc9dn electroporated embryo (C). (H) and (I) are higher magnifications of the posterior limb mesenchyme on the contralateral (H) and GFP/Hoxc9dn electroporated (I) sides. Note the significant decrease in Shh expression in the electroporated limb mesenchyme despite the mosaicism in expression due to the electroporation technique. (J) Measurements of the area of Shh expression domain of GFP-(n=10 embryos) and GFP/Hoxc9dn- (n=13 embryos) electroporated limbs, normalized to the contralateral side. Error bars represent SEM. Mann-Whitney statistical test, *** p<0,001. Scale bar is 100μm.

## Supplemental Movie Legends

### Movie S1. Related to Figure 2. Progressive formation of the LPM during gastrulation

17h time-lapse movie of a chicken embryo electroporated with cytoplasmic GFP and nuclear H2b-RFP fluorescent reporters, showing the dynamic process of LPM formation (from stage 5 to stage 11). Images were acquired on an inverted 2-photon microscope, using a long distance 10x objective. White bracket in the 1^st^ frame outlines the presumptive LPM (prLPM); white asterisk represents the Hensen node; white dashed line outlines the primitive streak (PS); white brackets in the last frame outline the forelimb (FL), interlimb (IL) and hindlimb (HL) domains.

### Movie S2. Related to Figure 2. Retrospective tracking of LPM precursors to their epiblast origin

Example of retrospective cell tracking of forelimb, interlimb and hindlimb LPM cells to identify their epiblast origin on a 24h time-lapse movie of a GFP and H2b-RFP-electroporated embryo. Images were acquired on an inverted 2-photon microscope, using a long distance 10x objective. Left panel, bright field image of the embryo (from stage 11 back to stage 4). Middle panel, GFP electroporated cells. Right panel, tracking of 9 LPM cells back to their epiblast origin. Note that the movie plays backwards.

### Movie S3. Related to Figure 2. Photoconverted mEOS2 transgenic embryos showing the LPM progressive formation

*hUbC:mEOS2FP* transgenic quail embryos in which the presumptive LPM in the primitive streak has been photoconverted. Embryos were further imaged every 2h to follow the fate of the photoconverted region. Photoconversion and acquisition were performed on an inverted confocal microscope. Left, middle-left, middle-right and right panels show embryos where the presumptive LPM was photoconverted at stage 6, 7, 9 and 10, respectively. Black arrowheads outline the anterior limit of photoconverted cells.

